# Super Learner Ensemble Modeling of CPTAC Proteomic Data for Survival Prediction in Head and Neck Squamous Cell Carcinoma

**DOI:** 10.64898/2026.06.11.731237

**Authors:** Ethan Park, Hannah Lee, Eun Jeong Oh, Tristan Tham, Seungjun Ahn

## Abstract

Survival analysis in head and neck squamous cell carcinoma (HNSCC) is traditionally performed using Cox proportional hazards models, alongside some exploration into black-box machine learning methods. The Super Learner (SL) algorithm addresses this model selection dilemma by combining diverse candidate algorithms into a weighted ensemble to perform comparably to the best candidate method. This study evaluates the performance of SL in HNSCC. Proteomic features as well as clinical covariates from 96 CPTAC HNSCC samples were modeled with three candidate algorithms (Cox LASSO, Cox Ridge, and Random Survival Forest) as well as the ensemble SL method. Models were optimized via Uno’s time-dependent Concordance Index (C-index) and tested at 1- and 3-year time horizons using 2000 bootstrap resamples. The Cox Ridge regression model achieved the highest predictive accuracy among the four total methods. However, the SL demonstrated stable performance over both time horizons (1-year C-index: 0.985; 3-year C-index: 0.960). Variable importance analysis of the Cox Ridge model successfully identified malignant proteins (ATR, MAML1, MIEN1) alongside novel potential prognostic indicators (ZNF800, KERA). This analysis emphasizes the statistical necessity for larger cohorts for ensemble learning, while providing a benchmark of proteomic indicators in HNSCC.

## Introduction

Head and neck squamous cell carcinoma (HNSCC) are head and neck cancers that arise from the mucosal epithelium in the oral cavity, pharynx, and larynx. It is the sixth most common cancer worldwide, with its incidence projected to increase by 30% by 2030, totaling 1.08 million new cases annually.^1^ Standard treatment of HNSCC is typically multimodal, including surgical resection, radiation, and chemotherapy with several novel treatments, including immunotherapies, in clinical trials.^2^ In recent years, studies have leveraged proteomic data to characterize molecular differences across HNSCC subtypes yielding prognostic insights and enabling the development of proteomic-based prognostic models in survival prediction.^3, 4, 5, 6^ However, survival analysis in HNSCC remains largely dominated by the Cox proportional hazards (PH) model to estimate the hazard ratios.^7^ There has been some exploration into black-box machine learning (ML) models such as random survival forest (RSF)^8^ including a recent study combining Cox PH and RSF analyses using SEER data.^9^ There is currently conflicting evidence on whether Cox-based models or ML techniques would be superior in predicting the prognosis of patients in HNSCC.^10,11^ The Cox model may not be valid when its main assumption, namely the PH assumption, is violated. While these ML models such as RSF have shown promising predictive performance, no single model has been shown to be universally optimal.^12, 13, 14^

The Super Learner (SL hereafter) algorithm^15^ offers a theoretical solution to the problem of model selection. SL was originally developed as an extension of stacked generalization, an ensemble learning framework that combines predictions from multiple candidate models.^16^ By creating an ensemble of multiple models and performing *k*-fold cross-validation to estimate the optimal weighted combination of these models, SL is asymptotically guaranteed (as *n*→∞) to perform at least as well as the best performing candidate model. In this asymptotic case, the SL may assign most or all of the weight to the best performing model.^17^ We hypothesize that a SL-based survival prediction model incorporating high-dimensional protein expression data along with clinical and demographic covariates will achieve higher predictive accuracy than individual candidate learners (i.e., LASSO- or SCAD-penalized Cox regression and RSF), as measured by the cross-validated concordance index (c-index).

The use of SL has mainly been applied in non-oncology settings, with limited applications in survival analysis.^18, 19, 20^ The literature lacks comprehensive evaluations of SL for oncology survival prediction, which motivates this present research. To our best knowledge, no prior studies have applied an SL ensemble to proteomic data for predicting survival outcomes in HNSCC, nor have they systemically evaluated its performance against individual candidate learners for HNSCC survival prediction using proteomic data. Furthermore, this study investigates the performance and practical limitations of SL when applied to high-dimensional, low-sample-size (HDLSS) settings. Given the limited sample sizes and data availability in many oncology datasets, this analysis is of particular importance.

## Methods

### Data Source and Study Cohort

The pre-processed proteomics (protein expression) and clinical data used in this study were obtained from the Clinical Proteomic Tumor Analysis Consortium (CPTAC) through the Proteomic Data Commons (https://pdc.cancer.gov/pdc/cptac-pancancer), which is one of the largest public repositories of proteogenomic data. The CPTAC is a National Cancer Institute (NCI)-driven initiative that provides large-scale mass spectrometry-based proteomics data, with greater protein coverage than the Cancer Genome Atlas (TCGA).^21^

Our dataset contains N = 96 samples with both protein expression and clinical data available. For each sample, 9,469 protein expression features were available. In addition to proteomic features, the dataset also includes nine clinical and demographic covariates as follows: sex, age, BMI, alcohol consumption history, tobacco smoking history, histologic grade, tumor size, tumor site curated, and cancer stage. A consolidated standards of reporting trials (CONSORT)-style diagram is provided in Figure 1.

**Figure 1.**
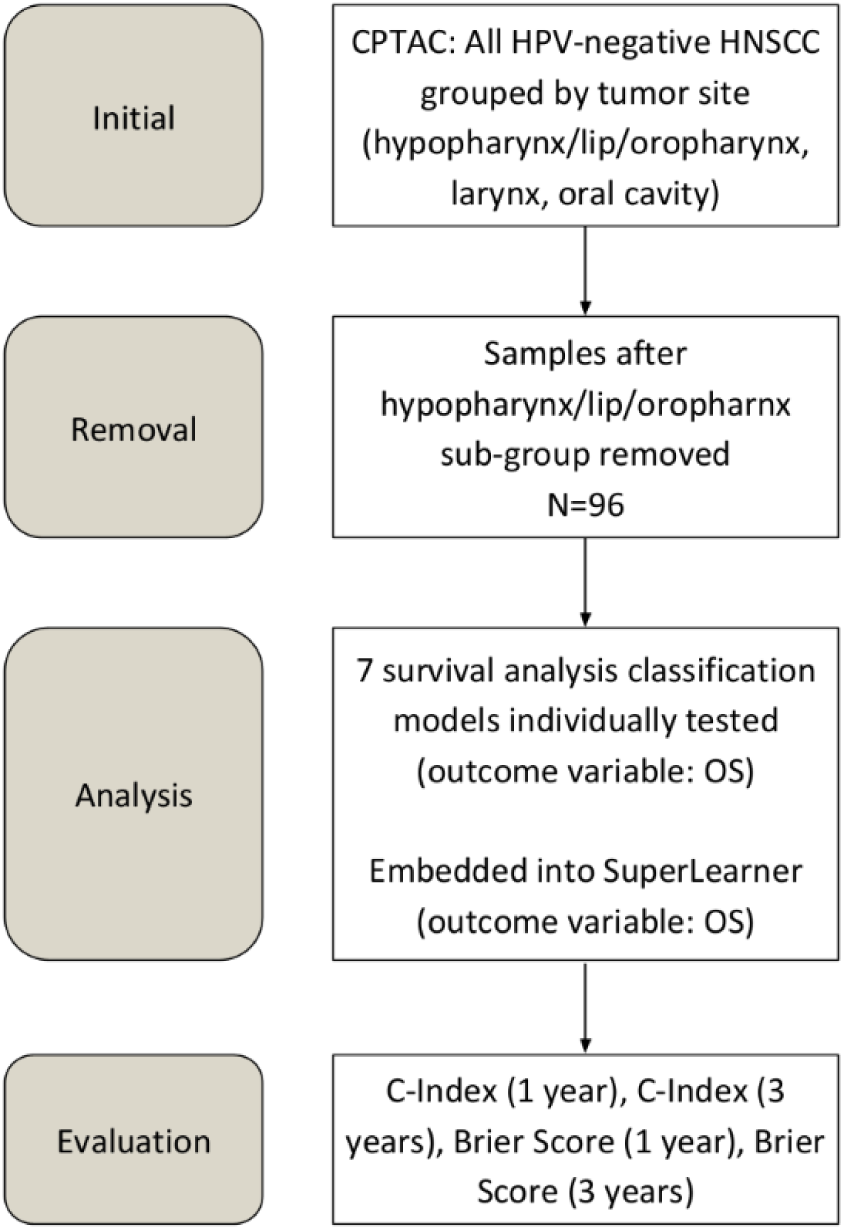
Data CONSORT Diagram. Dataset: CPTAC HNSCC Predictor variable: protein expression levels along with clinical covariates Outcome variable: Overall survival (OS), defined as the time elapsed before death of any cause Model Training/Testing: Trained on full dataset, testing consisted of 2000 bootstrapped samples (random resampling method with replacement, each subpopulation size = 96)

### Study Variables

The outcome variable was overall survival (OS), defined as the time from the first diagnosis of the disease to death from any cause, with patients censored at the date of last contact if alive. Predictor variables included protein expression features and the aforementioned nine clinical and demographic covariates (see Data Source and Study Cohort subsection above). BMI was categorized based on the following cutoffs: underweight (< 18.5 kg/m^2^), normal weight (18.5 to 25 kg/m^2^), overweight (25 to 30 kg/m^2^), and obese (> 30 kg/m^2^). Patients whose tumor sites were labeled under the group of hypopharynx and lip, were excluded because the management and prognosis of these cancers is substantially different and also because of the small sample sizes.^22^ Additionally, oropharynx was chosen to be removed because HPV status, a significant prognostic factor, was not available in the database.^23^

### Statistical Analysis

All statistical analyses were performed using R statistical software version 4.4.3 (R Foundation for Statistical Computing, Vienna, Austria).

### Model Specification

Using the *survivalSL* package version 0.98, we fit a SL model to the complete analysis dataset using the following candidate learners: LASSO-penalized Cox regression (LIB_COXlasso), ridge-penalized Cox regression (LIB_COXridge), and RSF (LIB_RSF). These learners were selected to represent commonly used regression-based and machine learning approaches for cancer survival prediction.

### Hyperparameter Tuning and Model Fitting Setting

Hyperparameters were tuned for the penalized Cox learners. For COX_lasso and COX_ridge, the selected optimal lambda values were 10 and 9.97, respectively. We allowed the SL to automatically estimate the hyperparameters for RSF, resulting in the following configurations: nodesize = 30; mtry = 4744; ntree = 500. The number of cross-validation folds was set to 5. The parameter optim.local.min was set to “TRUE” to ensure the numerical optimization. We assigned a penalty factor of 0 to all clinical covariates, as these variables are typically considered important confounders to adjust for and were therefore left unpenalized. The metric parameter was set to Uno’s c-index^24^, so that model weights were estimated using Uno’s c-index as the optimization criterion.

### Model Validation and Performance Evaluation

Due to the limited sample size (N = 96), we trained the model on the entire dataset, and model performance was assessed using bootstrap resampling rather than a separate test set.^25^ We generated 2000 bootstrap resamples, each of the same size as the original dataset, by sampling with replacement. For each bootstrap resample, time-dependent Uno’s c-index and the time-dependent Brier score at 1- and 3-year prediction horizons were computed. For the SL ensemble and each candidate learner, survival predictions were obtained as the estimated probability of surviving beyond the specified time horizon. Metrics were computed using the built-in *metrics* function in *survivalSL* package with *metric = “ci”* for the time-dependent c-index and *metric* = “*bs*” for the time-dependent Brier score. Mean estimates and 95% confidence intervals (CI) were reported across bootstrap resamples.

Given the censoring rate in our dataset (33%), Uno’s c-index was used as the primary concordance metric because it accounts for censoring through inverse probability of censoring weighting (IPCW).^26, 27^ This approach was selected to reduce the censoring sensitivity associated with Harrell’s c-index.^28^

### Variable Importance Analysis

After performance evaluation, we used the best performing individual learner on the complete analytic dataset to compute variable importance (VIMP) scores and identify proteins with the greatest contributions to survival prediction.

## Results

*Table 1* summarizes the baseline statistics of our study data, stratified by censoring status. The p-value for each covariate with censoring outcomes was also calculated. The study population primarily consisted of older adults, with a median age of 62.0 years. The cohort was overwhelmingly male (n = 84, 87.5%). More than half of the patients were of normal weight (n = 53, 55.2%), followed by overweight (n = 22, 22.9%) and obese (n = 15, 15.6%).

**Table 1.**
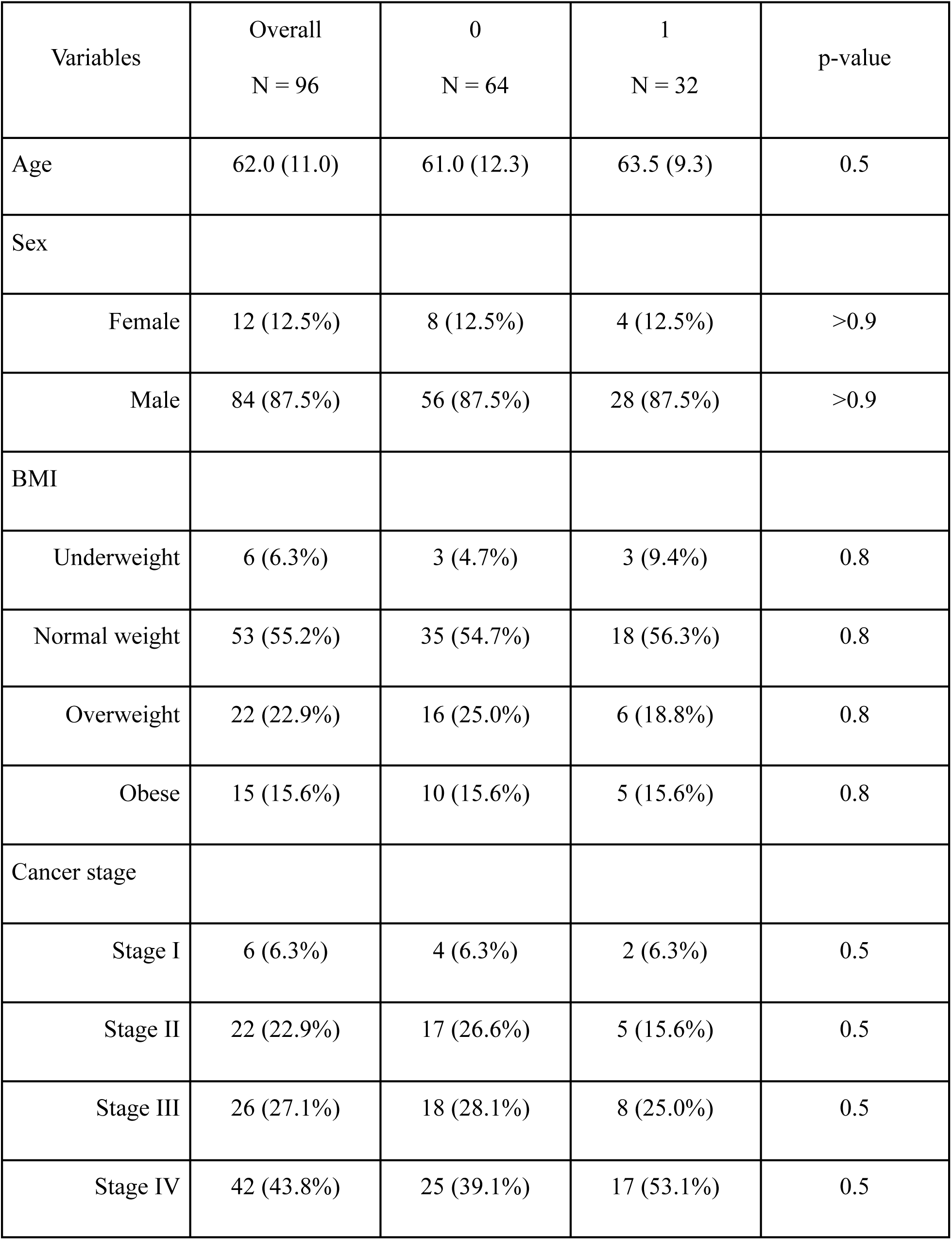
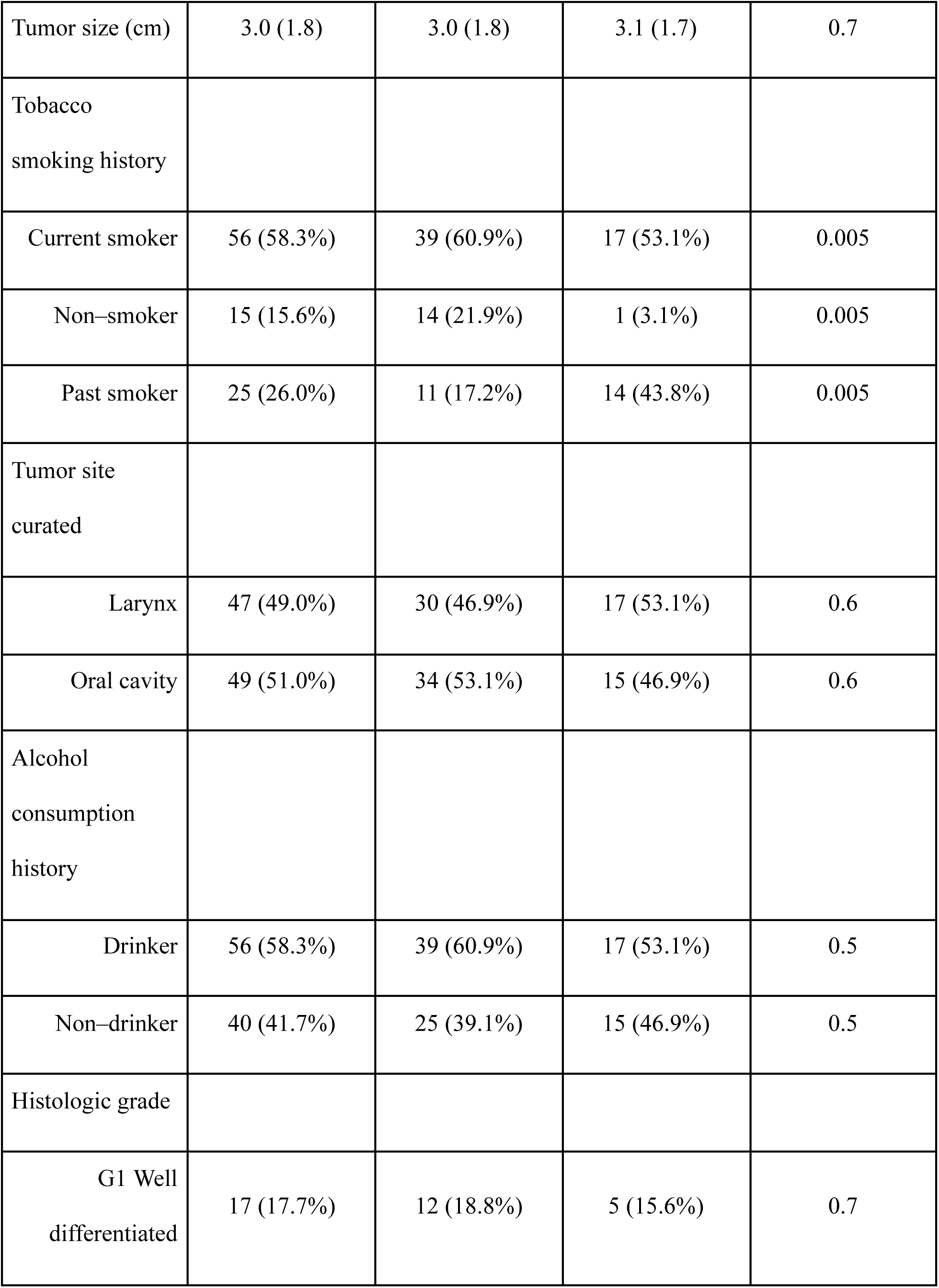
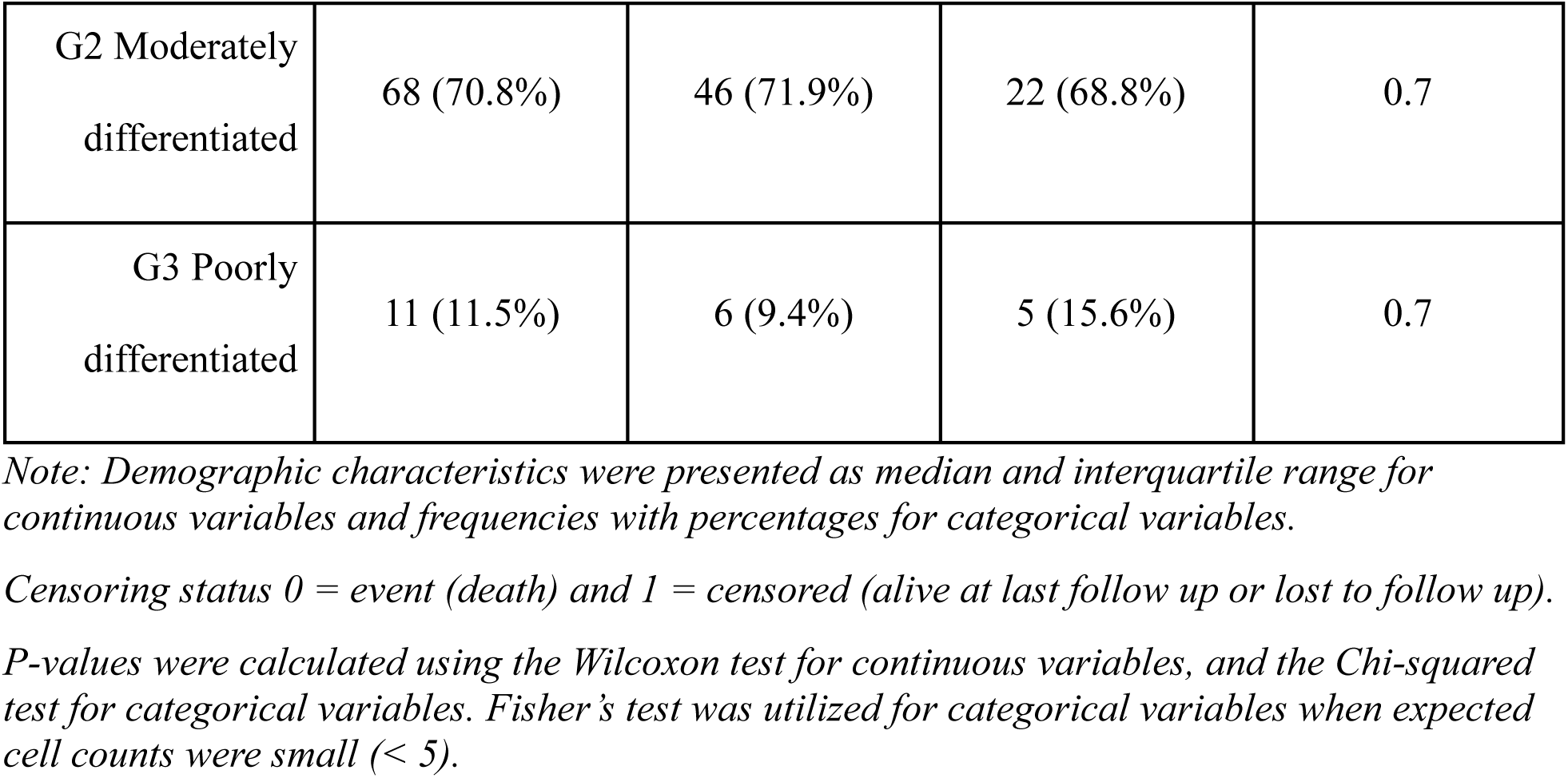
Descriptive statistics of study cohort stratified by censoring status.

| Variables | Overall<br>N = 96 | 0<br>N = 64 | 1<br>N = 32 | p-value |
| --- | --- | --- | --- | --- |
| Age | 62.0 (11.0) | 61.0 (12.3) | 63.5 (9.3) | 0.5 |
| Sex |  |  |  |  |
| Female | 12 (12.5%) | 8 (12.5%) | 4 (12.5%) | >0.9 |
| Male | 84 (87.5%) | 56 (87.5%) | 28 (87.5%) | >0.9 |
| BMI |  |  |  |  |
| Underweight | 6 (6.3%) | 3 (4.7%) | 3 (9.4%) | 0.8 |
| Normal weight | 53 (55.2%) | 35 (54.7%) | 18 (56.3%) | 0.8 |
| Overweight | 22 (22.9%) | 16 (25.0%) | 6 (18.8%) | 0.8 |
| Obese | 15 (15.6%) | 10 (15.6%) | 5 (15.6%) | 0.8 |
| Cancer stage |  |  |  |  |
| Stage I | 6 (6.3%) | 4 (6.3%) | 2 (6.3%) | 0.5 |
| Stage II | 22 (22.9%) | 17 (26.6%) | 5 (15.6%) | 0.5 |
| Stage III | 26 (27.1%) | 18 (28.1%) | 8 (25.0%) | 0.5 |
| Stage IV | 42 (43.8%) | 25 (39.1%) | 17 (53.1%) | 0.5 |
| Tumor size (cm) | 3.0 (1.8) | 3.0 (1.8) | 3.1 (1.7) | 0.7 |
| Tobacco<br>smoking history |  |  |  |  |
| Current smoker | 56 (58.3%) | 39 (60.9%) | 17 (53.1%) | 0.005 |
| Non-smoker | 15 (15.6%) | 14 (21.9%) | 1 (3.1%) | 0.005 |
| Past smoker | 25 (26.0%) | 11 (17.2%) | 14 (43.8%) | 0.005 |
| Tumor site<br>curated |  |  |  |  |
| Larynx | 47 (49.0%) | 30 (46.9%) | 17 (53.1%) | 0.6 |
| Oral cavity | 49 (51.0%) | 34 (53.1%) | 15 (46.9%) | 0.6 |
| Alcohol<br>consumption<br>history |  |  |  |  |
| Drinker | 56 (58.3%) | 39 (60.9%) | 17 (53.1%) | 0.5 |
| Non-drinker | 40 (41.7%) | 25 (39.1%) | 15 (46.9%) | 0.5 |
| Histologic grade |  |  |  |  |
| G1 Well<br>differentiated | 17 (17.7%) | 12 (18.8%) | 5 (15.6%) | 0.7 |
| G2 Moderately differentiated | 68 (70.8%) | 46 (71.9%) | 22 (68.8%) | 0.7 |
| G3 Poorly differentiated | 11 (11.5%) | 6 (9.4%) | 5 (15.6%) | 0.7 |

*Note: Demographic characteristics were presented as median and interquartile range for continuous variables and frequencies with percentages for categorical variables.*
*Censoring status 0 = event (death) and 1 = censored (alive at last follow up or lost to follow up).*
*P-values were calculated using the Wilcoxon test for continuous variables, and the Chi-squared test for categorical variables. Fisher's test was utilized for categorical variables when expected cell counts were small (< 5).*

*Table 2* summarizes performance metrics for each individual learner, as well as the SL. Ridge-penalized Cox regression model showed the best performance across all four metrics (i.e., 1- and 3-year Brier scores and 1- and 3-year Uno’s c-indices) for overall survival prediction in HNSCC. Although the SL did not outperform ridge-penalized Cox model, it showed strong predictive performance, with a 1-year c-index of 0.985 (95% CI: 0.962, 1.000). These findings suggest that high-dimensional proteomic data, combined with clinical covariates, can provide strong predictive signals for OS.

**Table 2.**
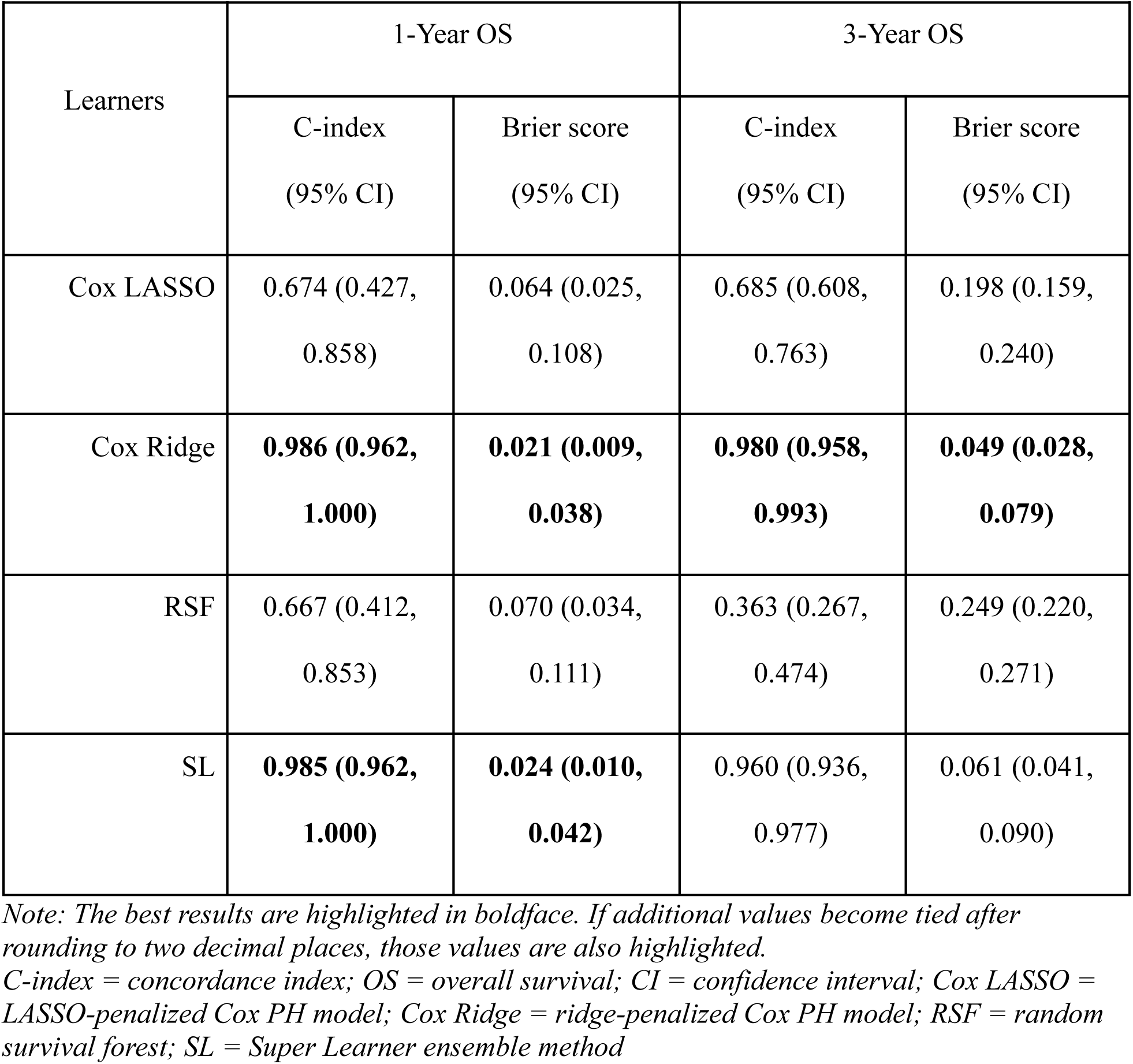
Summary of performance metrics for survival prediction models across 2000 bootstrap resamples.

*Table 3* displays the estimated weights for each learner, obtained through 5-fold cross-validation. The ridge-penalized Cox model received the largest ensemble weight, which is consistent with its highest predictive accuracy shown in Table 2.

**Table 3.** Super Learner weights for each individual candidate learner.

|  |  |
| --- | --- |
| Cox LASSO | 0.06925 |
| Cox Ridge | 0.81751 |
| RSF | 0.11322 |

*Table 4* presents the results of the ridge-penalized Cox regression analysis, which performed the best in predicting OS. The 10 proteins with the highest VIMP scores were extracted. A higher VIMP score indicates that the model relied more heavily on the corresponding protein for survival prediction. Clinically, proteins with high VIMP scores may be considered potentially important prognostic markers in HNSCC, although these findings should be carefully interpreted as model-based associations rather than causal effects. A positive coefficient indicates higher risk, in other words, higher expression of the protein is associated with shorter survival time. A negative coefficient indicates a protective association, meaning that higher expression of the protein is associated with longer survival time. The 10 most contributory proteins were ZNF800, KERA, ATR, MIEN1, POLR2E, MIEF1, MAML1, PARS2, ZNF609, and AP4S1. Higher expression of KERA, MIEN1, POLR2E, PARS2, and ZNF609 was associated with worse survival, whereas higher expression of ZNF800, ATR, MIEF1, MAML1, and AP4S1 was associated with longer survival.

**Table 4.** Top 10 proteins by VIMP scores from the Cox Ridge analysis.

| Protein (Gene Symbol)* | Coefficient | Importance Score |
| --- | --- | --- |
| ZNF800 | -0.008537109 | 0.008537109 |
| KERA | 0.008344348 | 0.008344348 |
| ATR | -0.008239223 | 0.008239223 |
| MIEN1 | 0.008098634 | 0.008098634 |
| POLR2E | 0.007728809 | 0.007728809 |
| MIEF1 | -0.007226214 | 0.007226214 |
| MAML1 | -0.007073833 | 0.007073833 |
| PARS2 | 0.007065508 | 0.007065508 |
| ZNF609 | 0.006998233 | 0.006998233 |
| AP4S1 | -0.006968741 | 0.006968741 |

\*Gene symbols are shown only as protein annotations and were mapped from Ensembl identifiers.

## Discussion

Our study findings are consistent with prior simulation studies, which have demonstrated that tree-based machine learning algorithms such as random forest do not perform well in high-dimensional data settings.^29, 30^ Multiple studies suggest that RSF may be better suited than Cox-based methods for modeling complex data; however, RSF generally requires a larger number of events for stable model training.^31^ This aligns with statistical theory regarding the efficacy of such learners in high-dimensional settings, particularly in proteomics data where feature collinearity is highly probable.^32^

The underperformance of the ensemble SL in comparison to the highest performing learner (ridge-penalized Cox model) is consistent with prior literature, with several studies outlining the potential limitations of using ensemble prediction models in HDLSS data. While the SL is asymptotically designed to outperform all individual models, we have concluded that ensemble models may underperform in finite-sample settings due to high variance in estimating ensemble weights when the sample size is limited.

Although the SL did not outperform all individual learners in this analysis, it showed relatively stable predictive performance across time horizons. While RSF showed moderate short-term predictive performance (1-year c-index: 0.667), its performance was substantially lower for long-term survival prediction (3-year c-index: 0.363). In contrast, the SL ensemble maintained high performance at both time horizons (1-year c-index: 0.985; 3-year c-index: 0.960). These findings suggest that the SL ensemble may have benefited from the penalized regression learners, which provided more stable predictions than RSF in this dataset. Overall, these results do not rule out the potential value of ensemble learning approaches for survival prediction. Further work is warranted to evaluate whether alternative learner libraries, tuning strategies, or larger datasets may improve SL performance.

Our analysis identified ZNF800, KERA, ATR, MIEN1, POLR2E, MIEF1, MAML1, PARS2, ZNF609, and AP4S1 as the 10 most contributory proteins to OS. Of these, ATR, MAML1, and MIEN1 are known contributors to HNSCC pathogenesis. ATR and MAML1 have well-established roles in HNSCC. ATR is a DNA damage signaling kinase that has been observed to be overexpressed in HNSCC and hypothesized to lead to reduced cytotoxic effects of conventional treatments such as chemotherapy and radiation as ATR activity may promote DNA repair within tumor cells.^33^ MAML1 is a transcriptional co-activator of the Notch signaling pathway, among other critical pathways such as Wnt/B-catenin, Sonic Hedgehog, and Hippo pathways. Several studies have demonstrated loss of function mutations in MAML1 leading to increased frequency of HNSCC tumor occurrence as well as poorer differentiation of tumors, correlating with its negative coefficient, or protective function, observed in our analysis.^34, 35, 36^ MIEN1, a gene associated with aiding cell migration and invasion, has been identified by Rajendiran et al. as being overexpressed in oral squamous cell carcinoma and its inhibition reducing cancer cell migration and invasion in vitro.^37^ In addition, their study demonstrated that higher MIEN1 expression correlated with poorer survival based on TCGA data, a finding present in our study’s analysis of the CPTAC data. The identification of previously reported prognostic proteins by the ridge-penalized Cox model supports the biological validity of the CPTAC database and its application in identifying potential biomarkers in HNSCC. Furthermore, as the ridge-penalized Cox model was included as a base learner in the SL ensemble, this suggests that SL has access to clinically relevant and meaningful predictors and supports its use in predicting survival outcomes.

Our results also showed that POLR2E played a contributory role in predicting survival outcomes. The POLR2E rs378016 polymorphism has been found to increase cancer risk in prostate, breast, cervical, thyroid, esophageal, and gastric cancer.^38, 39, 40^ Though no specific studies have identified POLR2E as a risk factor in HNSCC, its prevalence in the pathogenesis of other malignancies suggests a conserved role that may extend to the development of HNSCC and may warrant future investigation. Similarly, there have been no specific studies demonstrating the role of ZNF800 or ZNF609 in the pathogenesis of HNSCC, though several members of the Zinc finger proteins (ZNF), including ZNF14, ZNF134, ZNF160, ZHF420, ZNF610, ZHF880, have been studied as potential biomarkers in HNSCC.^41, 42, 43^ Overexpression of MIEF1 was found in non-small cell lung cancer and invasive breast carcinoma; however, its role in HNSCC has not been studied.^44^ PARS2, APS1, and KERA have not been associated with cancer biology, nor have they been identified as potential cancer biomarkers and may represent either novel biological findings or model artifacts and should thus be cautiously interpreted.

Due to the constraints of HNSCC dataset sample size which restricts the use of a full train-test split, our proteomic review is restricted to the most influential proteins identified by the ridge-penalized Cox model. This analysis provides a baseline for the proteins most strongly associated with HNSCC prognosis.

## Data Availability

The datasets analyzed for this study are publicly available in the Clinical Proteomic Tumor Analysis Consortium (CPTAC) repository though the Proteomic Data Commons at https://pdc.cancer.gov/pdc/cptac-pancancer.

